# Reticulation-Aware Dissimilarities for Phylogenetic Networks

**DOI:** 10.64898/2026.11.08.765198

**Authors:** Ziyun Guang, Mark Kessler, Nicolae Sapoval, Zhi Yan, Luay Nakhleh

## Abstract

Dissimilarity measures play a critical role in phylogenetics. They help quantify the accuracy of inferences when the ground truth is known, allow comparisons of inferences obtained using different methods, and provide a tool for summarizing sets of phylogenies. A wide array of dissimilarity measures has been introduced for phylogenetic trees over the past half-century, and they vary in their mathematical properties, empirical behavior, and computational complexity. When the evolutionary history involves reticulate processes, such as hybridization and horizontal gene transfer, it is represented by phylogenetic networks. Although the study of phylogenetic network dissimilarity is less established than its tree counterpart, the past two decades have seen an increasing number of measures developed for phylogenetic networks. However, while some have established mathematical properties—for example, metricity on certain classes of phylogenetic networks—none properly separates reticulation disagreement from tree-like disagreement. This is the rationale behind the introduction of the hybrid ancestry error (HAE) measure by Dai and Molloy. The HAE measure, however, is too simplistic to distinguish different reticulation scenarios. In this work, we propose a novel family of network dissimilarity measures that explicitly account for reticulation differences while being more informative than HAE. We prove that our measures are pseudometrics on rooted binary phylogenetic networks on a fixed taxon set, with two instantiations yielding metrics on binary level-1 networks. Empirically, the measures remain sensitive to reticulation errors as network size grows and respond to changes in hybrid identity and parental placement. Calibration experiments further characterize the measures’ behavior under network edits of various magnitudes.

## 1 Introduction

Dissimilarity measures are essential in the field of phylogenetics for evaluating inferred evolutionary histories, comparing results across inference methods, and summarizing collections of phylogenies. Over the past half-century, the field has established a wide array of measures for phylogenetic trees, including the Robinson–Foulds distance [23] and rearrangement distances [1]. These measures differ in their mathematical properties, sensitivity to structural changes, and computational complexity. Trees alone, however, cannot represent reticulate evolutionary histories involving hybridization, introgression, and horizontal gene transfer [6, 16]. Representing these histories requires phylogenetic networks [9].

Methods have also been proposed for phylogenetic network comparisons, typically focusing on their underlying graph topology. These approaches describe the topology through hardwired or softwired clusters [9], path multiplicities [4], recursive vertex structure [21], or edge-based tripartitions [22, 18]. Several are metrics on particular classes of phylogenetic networks. However, their overall scores do not explicitly separate reticulation disagreement from disagreement in the tree-like part of the network. In particular, normalized measures based on global structural summaries can assign small dissimilarities despite substantial reticulation errors when the tree-like portion of the network is extensive.

As inference methods scale to larger numbers of taxa, this distinction becomes particularly important. To address this, Dai and Molloy [5] recently introduced the hybrid ancestry error (HAE), which compares the sets of taxa descending from at least one reticulation in the reference and inferred networks. They reported that end-to-end methods could achieve low or moderate global topology error, measured by the normalized hardwired-cluster distance [9], while recovering hybrid ancestry poorly. Overall topological agreement therefore need not imply accurate recovery of reticulate ancestry. This matters because recovering these reticulate relationships is a central goal of network inference. A missing reticulation overlooks a potential exchange of genetic material, whereas an incorrectly placed reticulation attributes that exchange to the wrong lineages.

However, HAE records only which taxa have hybrid ancestry; it does not distinguish the reticulation events responsible or their parental lineages. Networks with different numbers or placements of reticulations can have zero HAE when the same taxa have hybrid ancestry. We therefore seek a dissimilarity that explicitly compares the number of reticulations and, for each reticulation, which hybrid lineage receives genetic material and which parental lineages contribute to it. It should retain sensitivity to missing and misplaced reticulations as the tree-like portion of the network grows. Disagreement in the remaining tree structure should remain visible. These requirements motivate a construction with separately interpretable reticulation and tree-remainder terms.

We introduce a family of network dissimilarities based on optimal matching of reticulation tripartitions^1^. Each tripartition represents the hybrid and parental subnetworks associated with one reticulation. The reticulation term, *D*_ret_, compares these representations through an optimal assignment, with unmatched reticulations accounting for differences in their numbers. Independently, the tree-remainder term, *D*_tree_, compares the remainders obtained by deleting reticulation vertices and their descendants. Marker leaves record where the removed subnetworks were attached. Their combination yields a joint dissimilarity while preserving the distinction between reticulation and tree-remainder disagreement.

We develop three instantiations based on leaf-set Jaccard, cluster-multiset Jaccard, and vertex-equivalence dissimilarities. We prove that our dissimilarities are pseudometrics on the space of rooted binary phylogenetic networks with the same taxon set. We also prove that the cluster and vertex-equivalence instantiations are metrics on binary level-1 networks. All resulting dissimilarity measures are computable in polynomial time with respect to the input network sizes. Empirically, through controlled perturbations and comparisons of inferred networks, we show that the measures remain sensitive to missing reticulations and distinguish changes in hybrid identity, parental placement, and tree structure. We show how explicitly comparing reticulations can complement overall topological assessments and help explain differences among inferred networks.

## 2 Background and notation

A *phylogenetic network N* on a taxon set *X* is a rooted directed acyclic graph with vertex set *V* (*N*) and edge set *E*(*N*). Its vertices comprise a unique *root* of indegree 0, *leaves* of outdegree 0 bijectively labelled by *X*, internal *reticulation vertices* of indegree greater than 1, and internal *tree vertices* of indegree 1. We consider *binary networks*: the root and internal tree vertices have outdegree 2, reticulations have indegree 2 and outdegree 1, and leaves have indegree 1 [8, Definition 3]. We exclude parallel edges. We denote by *R*(*N*) the set of reticulation nodes of *N* and by *r*(*N*) = |*R*(*N*)| the number of reticulations.

A vertex *u* is an *ancestor* of *v* if there is a directed path from *u* to *v*. Let ℒ (*v*) *⊆ X* denote the set of leaves reachable from *v* by a directed path—this is the *hardwired cluster*, or *leaf-descendant set*, of *v*. The *cluster multiset* of a network contains one cluster per vertex, retaining repeated occurrences [8]. For a vertex *x* of *N* , the *subnetwork rooted at x*, denoted *N* (*x*), is the subgraph induced by *x* and its descendants. All vertices are retained without suppression. Its leaf set is *ℒ* (*x*). For a rooted subnetwork *S*, we also write *ℒ* (*S*) for its set of leaf labels.

Two networks *N*_1_ = (*V*_1_, *E*_1_) and *N*_2_ = (*V*_2_, *E*_2_) on the same taxon set *X* are *isomorphic*, written *N*_1_ *≅ N*_2_, if there exists a bijection *ϕ* : *V*_1_ *→ V*_2_ such that (*u, v*) *∈ E*_1_ ⇔ (*ϕ*(*u*), *ϕ*(*v*)) *∈ E*_2_, and *ϕ* maps every leaf to a leaf with the same label [3].

The *level* of a network is the maximum number of reticulations in any biconnected component of its underlying undirected graph. A network is *tree-child* if every non-leaf vertex has a child of indegree 1 [20]. Every binary level-1 network is tree-child [8, Proposition 9].

A *displayed tree* on *X* can be obtained by retaining one incoming edge at each reticulation, removing vertices and edges that no longer lead to a labelled leaf, and suppressing vertices of indegree 1 and outdegree 1, as well as a root of outdegree 1 when necessary [20]. A *tree of blobs* summarizes the tree-like relationships among reticulated regions by contracting each blob to a single vertex while preserving its connections to the rest of the network [5].

## 3 Tripartition dissimilarity measure

### Reticulation tripartition and dissimilarity

Deleting a vertex *h* from *N* removes *h* and all its incident edges [20, Def. 2]. We denote the resulting directed acyclic graph by *N ™ h*.

For a reticulation *h* with parents *u* and *v*, consider the subnetworks rooted at *u* and *v* in *N ™ h*. In each subnetwork, retain only vertices on directed paths from its root to an original labelled leaf of *N* . If no such path exists, the rooted subnetwork is defined to be empty.

Unlike the previously introduced edge-induced tripartitions of the taxon set [22, 18], our reticulation tripartitions associate each reticulation with three subnetworks, whose leaf sets may overlap.

#### Definition 1

(Reticulation tripartition). Let *h* be a reticulation node of *N* with parents *u* and *v*. The *reticulation tripartition* of *h* is the ordered triple

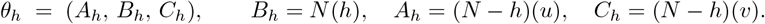

The middle block *B*_*h*_ contains *h* and all its descendants. The outer blocks *A*_*h*_ and *C*_*h*_ are the substructures rooted at the two parents once the hybrid is removed, retaining only vertices that lead to an original leaf node.

Hereafter, *tripartition* refers to this construction, and each of *A*_*h*_, *B*_*h*_, *C*_*h*_ is a *block*. The blocks may overlap. For the phylogenetic network comparisons below, we also allow empty networks, repeated leaf labels, roots of outdegree one, and vertices of indegree 1 and outdegree 1. Let *N* denote this extended class. Fix a dissimilarity *d* : *N × N →* [0, 1], defined also for networks with different taxon sets, with *d*(∅, ∅) = 0. We define the dissimilarity between two reticulation tripartitions as follows.

#### Definition 2

(Tripartition dissimilarity). Given two reticulation tripartitions *θ* = (*A, B, C*) and *θ*^*′*^ = (*D, E, F*), let

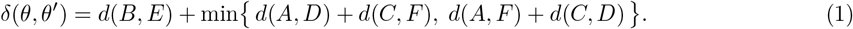

The middle blocks *B, E* are compared directly, while the parental blocks are compared under whichever of the two parent-to-parent correspondences is cheaper. Thus, *δ* is invariant under the swap (*A, B, C*) ↦ (*C, B, A*).

Let *N*_1_, *N*_2_ be networks on *X*. Their reticulation tripartitions form the multisets

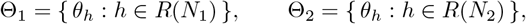

Each reticulation contributes one entry, including when different reticulations have identical tripartitions. Thus, |Θ_*i*_| = *r*(*N*_*i*_).

We match reticulations one-to-one. Let *m* = max *r*(*N*_1_), *r*(*N*_2_) , and pad the smaller tripartition multiset with dummy entries ⊥ until both multisets have size *m*. Since each block dissimilarity lies in [0, 1], the tripartition dissimilarity is at most 3. We therefore set the unmatched penalty to *ρ* = 3 and define

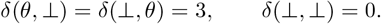

#### Definition 3

(Reticulation dissimilarity). Let Θ_1_ and Θ_2_ be the tripartition collections after padding if necessary. The *reticulation dissimilarity* of *N*_1_ and *N*_2_ is

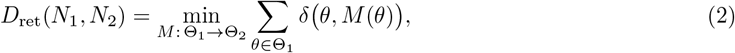

the minimum taken over all bijections *M* : Θ_1_ *→* Θ_2_. When both networks are trees, both tripartition collections are empty and we set *D*_ret_(*N*_1_, *N*_2_) = 0.

Each matching contains *m* pairs, each contributing at most 3. Define the normalized reticulation dissimilarity by

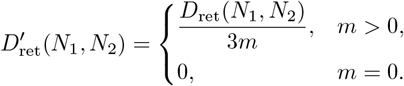

Thus, 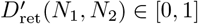.

### 3.2 Joint dissimilarity from reticulation tripartitions and a tree remainder

The reticulation dissimilarity *D*_ret_ does not capture all differences in the tree-like portion of the network. Two networks may have identical reticulation tripartitions but differ elsewhere. We therefore complement *D*_ret_ with a dissimilarity between their tree remainders, as illustrated in Figure 1.

**Figure 1.**
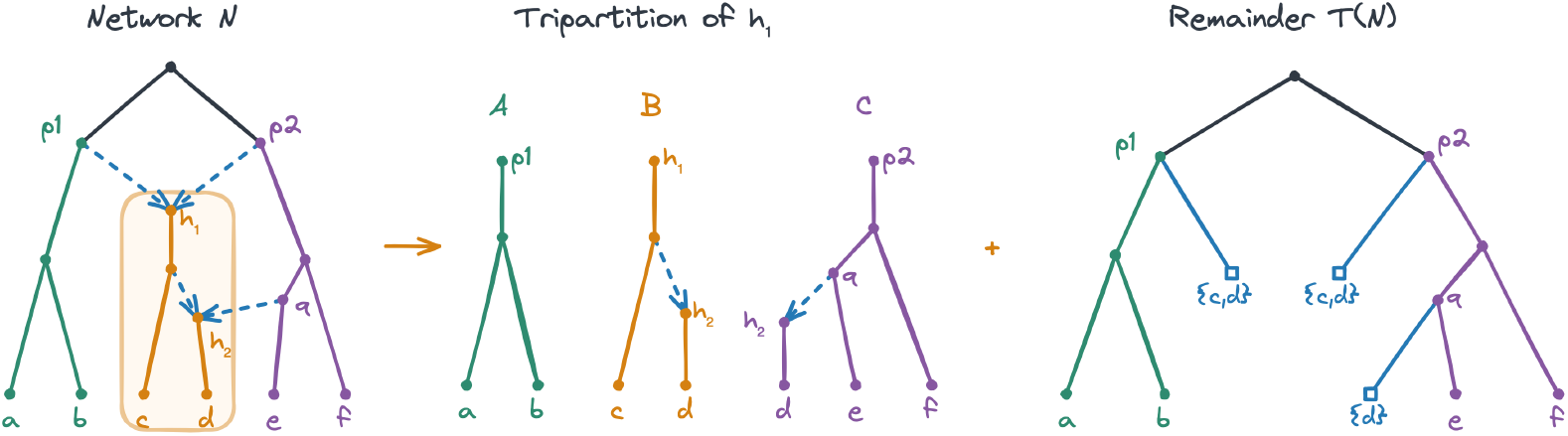
Illustration of the reticulation tripartition and the tree remainder. The middle block *B* contains *h*_1_ and its descendants; *A* and *C* are rooted at its parents after deleting *h*_1_. The vertex *h*_2_ remains in both *B* and *C*. In the marked remainder *T* (*N*), the two surviving parents of *h*_1_ each receive a marker labelled *{c, d}*. The surviving parent of *h*_2_ receives a marker labelled *{d}*.

#### The tree remainder

To obtain the tree remainder of network *N* , delete the union of the vertex sets of all middle blocks *B*_*h*_. For each original edge (*p, h*) whose parent *p* survives, attach a marker leaf to *p*. Label this marker by the original descendant taxon set ℒ (*h*), represented as a single atomic label distinct from taxon labels. Equal descendant taxon sets give equal marker labels across all networks. Retain all marker vertices, including those with repeated labels, and all surviving vertices without suppression. Write *T* (*N*) for the resulting marked remainder.

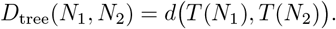

##### Definition 4

(Joint dissimilarity). The joint dissimilarity between *N*_1_ and *N*_2_ is

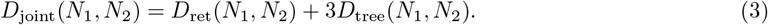

The factor 3 gives the tree-remainder term the same maximum contribution as one reticulation comparison. The normalized joint dissimilarity is

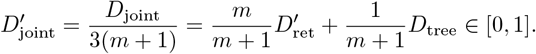

This convex combination assigns increasing weight to the reticulation term as *m* grows, which emphasizes reticulation recovery.

### 3.3 Instantiations of *d*

We consider three choices of *d*: leaf-set *d*_l_, cluster-multiset *d*_c_, and vertex-equivalence dissimilarity *d*_n_. We distinguish the resulting reticulation, remainder, and joint dissimilarities by the superscripts l, c, and n, respectively.

#### Leaf-set dissimilarity

For two subnetworks *S, S*^*′*^, let *P* = ℒ (*S*) and *Q* = ℒ (*S*^*′*^). Their leaf-set dissimilarity is the Jaccard distance

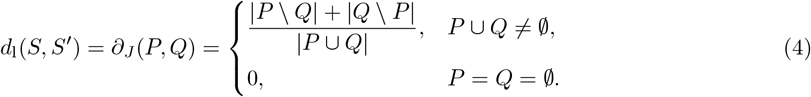

Repeated labels appear only once in these sets. This distance takes values in [0, 1]: it is 0 when the leaf sets coincide and 1 when they are disjoint and at least one is nonempty. Jaccard distance is a metric on finite sets [15, 17].

#### Cluster multiset dissimilarity

Unlike leaf-set dissimilarity, cluster-multiset dissimilarity compares internal structure through hardwired clusters [23, 9]. For a subnetwork *S*, let *C*(*S*) contain one hardwired cluster per vertex, retaining repeated clusters [8].

For a marked remainder, each cluster is itself a multiset containing one label per reachable leaf vertex; distinct marker leaves with equal labels contribute separate occurrences.

Define the cluster-multiset dissimilarity by

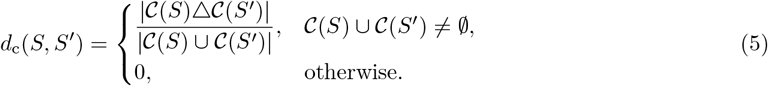

Here, multiset symmetric difference takes the absolute difference of occurrence counts, and multiset union takes their maximum.

We compare the full subnetworks without restricting them to shared leaf labels. The cluster multisets retain all vertices and directly account for differences in leaf labels, so no separate leaf-set penalty is needed. Substituting *d*_c_ into the generic definitions gives 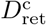 and 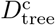.

#### Vertex-equivalence dissimilarity

Nakhleh [21] proposed a dissimilarity based on recursive representations of vertex structure, which we refer to here as the *vertex-equivalence dissimilarity* (VED). Two vertices are *equivalent* if they are leaves with the same label, or they are both internal nodes whose children admit a one-to-one correspondence in which corresponding children are equivalent.

Although originally defined for networks on the same taxon set, the vertex-equivalence formula also allows us to compare subnetworks with different leaf sets. An equivalence class absent from a subnetwork has multiplicity zero.

For each vertex-equivalence class *q*, let *c*_*S*_(*q*) be its multiplicity in *S*. Define

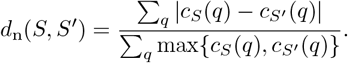

The sums range over all vertex-equivalence classes occurring in either subnetwork. For normalization, the denominator is the size of their multiset union. We use a multiset Jaccard normalization so that the block dissimilarity satisfies the triangle inequality. This differs from the total-vertex normalization used for the VED comparator in our experiments (Appendix C). Set *d*_n_(∅, ∅) = 0.

This is the Jaccard distance between the multisets of vertex-equivalence classes and therefore lies in [0, 1]. Nonempty blocks with disjoint leaf sets have dissimilarity 1.

### 3.4 Algorithm for computing the dissimilarities

We first construct the reticulation tripartitions and pad the smaller collection with dummy entries. We then compute the pairwise tripartition costs and solve the resulting minimum-weight perfect matching problem using the Hungarian algorithm [14, 19] — we construct the complete bipartite graph on the two collections of reticulation tripartitions Θ_1_ and Θ_2_, assigning weight *δ*(*θ, θ*^*′*^) to each pair based on their tripartition dissimilarity. Once the costs have been computed, the assignment step takes *O*(*m*^3^) time, where *m* = max {*r*(*N*_1_), *r*(*N*_2_)}. Independently, we construct and compare the marked tree remainders to obtain *D*_tree_ and the joint dissimilarity. Algorithm 1 in Appendix B summarizes the computation.

## 4 Theoretical Analysis

We first prove that all three joint dissimilarities are pseudometrics. We then prove that the cluster and vertex-equivalence instantiations are metrics on rooted binary level-1 networks. These results apply to the unnormalized joint dissimilarities.

### 4.1 Pseudometricity of the three instantiations

A metric on a set *S* is a function *d* : *S × S →* ℝ satisfying, for all *x, y, z ∈ S*:

i. **Nonnegativity:** *d*(*x, y*) *≥* 0.
ii. **Identity of indiscernibles:** *d*(*x, y*) = 0 if and only if *x* = *y*.
iii. **Symmetry:** *d*(*x, y*) = *d*(*y, x*).
iv. **Triangle inequality:** *d*(*x, z*) *≤ d*(*x, y*) + *d*(*y, z*).

A pseudometric retains nonnegativity, symmetry, the triangle inequality, and *d*(*x, x*) = 0, but permits *d*(*x, y*) = 0 for distinct *x, y*.

#### Lemma 1

(Triangle inequality for block and tripartition dissimilarities). *For each of the leaf-set, cluster, and vertex-equivalence instantiations, d and δ satisfy the triangle inequality*.

*Proof*. The leaf-set instantiation is the Jaccard distance on leaf-label sets. The cluster and vertex-equivalence instantiations are the Jaccard distances on the multisets of clusters and vertex-equivalence classes, respectively. The triangle inequality for Jaccard distance extends to multisets, as explicitly noted by Kosub [13]. Thus, *d* satisfies the triangle inequality in all three cases. Consider three reticulation tripartitions *θ*_*i*_ = (*A*_*i*_, *B*_*i*_, *C*_*i*_), for *i* = 1, 2, 3. Relabel the parental blocks so that optimal pairings between *θ*_1_, *θ*_2_ and between *θ*_2_, *θ*_3_ match the *A* blocks together and the *C* blocks together. Composing these pairings gives a candidate pairing between *θ*_1_ and *θ*_3_. By the triangle inequality for *d*,

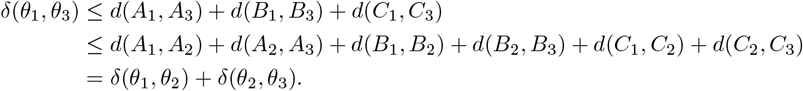

The triangle inequality also holds when dummy entries are present: real–real costs lie in [0, 3], real–dummy costs equal 3, and dummy–dummy costs equal 0. □

#### Lemma 2

(Pseudometricity of the reticulation dissimilarity). *For each α ∈ {*l, c, n*}*, 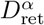 *is a pseudometric on rooted binary networks on a fixed taxon set*.

*Proof*. Fix one of the three instantiations. Nonnegativity and symmetry follow from the tripartition dissimilarity. Matching each reticulation to itself gives *D*_ret_(*N, N*) = 0. Now we prove the triangle inequality.

Let *N*_1_, *N*_2_, *N*_3_ be rooted binary networks on the same taxon set. Let *M*_12_ and *M*_23_ be minimizing matchings for *D*_ret_(*N*_1_, *N*_2_) and *D*_ret_(*N*_2_, *N*_3_), respectively.

Extend these matchings, if necessary, by adding dummy–dummy pairs of cost zero so that all three tripartition collections have cardinality max*{r*(*N*_1_), *r*(*N*_2_), *r*(*N*_3_)*}*, represented by Θ_1_, Θ_2_, Θ_3_. Their composition

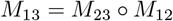

is a bijection from Θ_1_ to Θ_3_. Any matching with extra dummy entries can be converted to one under the original padding rule without increasing its total dissimilarity. Replace pairs (*θ, ⊥*) and (*⊥, θ*^*′*^) by (*θ, θ*^*′*^) and (*⊥, ⊥*), reducing their combined dissimilarity from 6 to at most 3. Repeat as needed and discard excess dummy–dummy pairs. By definition, *D*_ret_(*N*_1_, *N*_3_) is the minimum matching cost under the original padding rule. Thus, *D*_ret_(*N*_1_, *N*_3_) is no greater than the total dissimilarity of *M*_13_.

Applying Lemma 1 to each corresponding triple of tripartitions and summing gives

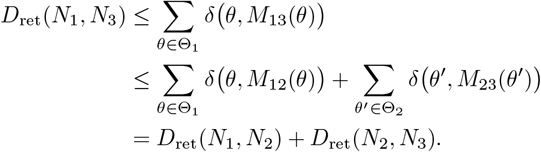

The second sum is reindexed using the bijection *M*_12_. Therefore, 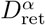 is a pseudometric. □

#### Lemma 3

(Pseudometricity of the tree remainder dissimilarity). *For each α ∈ {*l, c, n*}*, 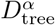 *is a pseudometric on rooted binary networks on a fixed taxon set*.

*Proof*. For each instantiation, *d* is nonnegative, symmetric, and zero when comparing a structure with itself. It satisfies the triangle inequality by Lemma 1. These properties also hold for our tree remainders, whose labels are determined independently for each network. The remainder comparisons are Jaccard distances on leaf-label sets, cluster multisets, or vertex-equivalence multisets, respectively. The triangle inequality holds for both set and multiset Jaccard distances, including repeated marker labels [13]. □

#### Theorem 1

(Pseudometricity of the joint dissimilarity). *For each α ∈ {*l, c, n*}*, 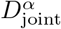 *is a pseudometric on rooted binary networks on a fixed taxon set*.

*Proof*. By Lemmas 2 and 3, 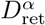 and 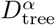 are pseudometrics. A positive weighted sum of pseudometrics is a pseudometric. Hence 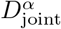 is a pseudometric on rooted binary networks on a fixed taxon set. □

### 4.2 Identity of indiscernibles and metricity

It remains to determine when zero joint dissimilarity implies isomorphism. We establish this property for the cluster and vertex-equivalence instantiations on rooted binary level-1 networks.

#### Claim 1.

*Let N be a rooted binary level-*1 *network and let h be a reticulation of N. The only edges entering B*_*h*_ *from outside are the two incoming edges of h, and no edge leaves B*_*h*_. *Consequently, every root-to-vertex path ending in B*_*h*_ *passes through h*.

*Proof*. In a binary level-1 network, every nontrivial biconnected component is an undirected cycle [8, Observation 16]. Recall that an edge is a bridge if and only if it belongs to no cycle in the underlying undirected graph. The two incoming edges of *h* lie on an undirected cycle, formed by two root-to-*h* paths through its distinct parents after their last common vertex before *h*. Thus, *h*’s unique outgoing edge (*h, w*) must be a bridge. Otherwise, a cycle containing (*h, w*) would also contain an incoming edge of *h* and hence belong to the same biconnected component as the first cycle. This biconnected component would contain all three edges incident to *h* and could not be a cycle.

Every directed path from *h* to a proper descendant starts with (*h, w*). Deleting this bridge therefore separates the root and *h* from all proper descendants of *h*. Hence every root-to-proper-descendant path passes through *h* and (*h, w*), ruling out any other incoming edge from outside *B*_*h*_ to a descendant of *h*. No edge leaves *B*_*h*_, since every child of a descendant of *h* is also a descendant of *h*. Consequently, every root-to-vertex path ending in *B*_*h*_ passes through *h*. □

Let *N* be a rooted binary level-1 network. Let ℋ be the set of reticulations of *N* having no proper reticulation ancestor. We prove the following lemma.

#### Lemma 4

(Decomposition of the removed vertex set for binary level-1 networks). *Blocks B*_*h*_, *h ∈* ℋ, *are pairwise vertex-disjoint, and* ∪_*h∈R*(*N*)_ *V* (*B*_*h*_) = ∪_*h∈*ℋ_ *V* (*B*_*h*_). *Thus, their vertex sets partition the vertex set removed when constructing the marked remainder*.

*Proof*. Suppose that *B*_*h*_ and *B*_*k*_ share a vertex *x* for distinct *h, k ∈ ℋ*. By Claim 1, every root-to-*x* path passes through both *h* and *k*. One must therefore be an ancestor of the other, contradicting the definition of *ℋ*. Thus, the blocks are pairwise vertex-disjoint.

Every reticulation *h* of *N* either belongs to *ℋ* or descends from some *k ∈ ℋ*, in which case *B*_*h*_ *⊆ B*_*k*_. Therefore, ∪_*h∈R*(*N*)_ *V* (*B*_*h*_) = ∪_*k∈ℋ*_ *V* (*B*_*k*_). This is precisely the vertex set removed when constructing the marked remainder. □

By Claim 1 and Lemma 4, a reticulation contributes markers to the remainder if and only if it belongs to *ℋ*; each such reticulation contributes exactly two markers.

#### Lemma 5

(Distinct descendant taxon sets of reticulations). *Let N be a rooted binary level-*1 *network. For distinct reticulations h, k of N*,

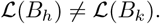

*Proof*. If neither reticulation is an ancestor of the other, their descendant taxon sets are disjoint: by Claim 1, a shared descendant leaf would place both reticulations on the same root-to-leaf path.

Otherwise, suppose that *h* is an ancestor of *k*. Then *ℒ* (*B*_*k*_) *⊆* ℒ (*B*_*h*_). Since *N* is tree-child, starting at *h* and repeatedly choosing a child of indegree 1 gives a path to a leaf that avoids *k*. Together with a root-to-*h* path, this gives a root-to-leaf path avoiding *k*. By Claim 1, this leaf is not a descendant of *k*. Thus, *ℒ* (*B*_*k*_) ⊊ *ℒ* (*B*_*h*_). In either case, the two descendant taxon sets differ. □

#### Lemma 6

(Reconstruction from the marked remainder and middle blocks). *Let N*_1_, *N*_2_ *be rooted binary level-*1 *networks on the same taxon set. Suppose there is a bijection M* : *R*(*N*_1_) *→ R*(*N*_2_) *such that*

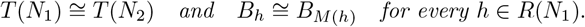

*Then N*_1_ *≅ N*_2_.

*Proof*. For *i* = 1, 2, let *ℋ*_*i*_ be the reticulations of *N*_*i*_ with no proper reticulation ancestor. By Lemma 4, the vertex sets of *B*_*h*_, *h ∈ ℋ*_*i*_, partition the vertices removed when constructing *T* (*N*_*i*_).

The block isomorphisms imply ℒ (*B*_*h*_) = ℒ (*B*_*M*(*h*)_). By Lemma 5, these taxon sets distinguish reticulations within each network. Since the remainder isomorphism preserves marker labels, it follows that *M* (*ℋ*_1_) = *ℋ*_2_ and that markers associated with *h* correspond to markers associated with *M* (*h*).

Reconstruct *N*_*i*_ by replacing the two markers associated with each *h ∈ ℋ*_*i*_ by edges from their parents to the root of *B*_*h*_. The remainder isomorphism maps corresponding marker parents to each other, and the block isomorphisms map corresponding roots to each other. Thus, the reconstruction preserves all attachment edges.

Since the removed blocks are pairwise vertex-disjoint, the remainder and block isomorphisms combine into an isomorphism *N*_1_ *≅ N*_2_. □

#### Theorem 2

(Identity of indiscernibles for the vertex-equivalence instantiation). *For rooted binary level-*1 *networks N*_1_, *N*_2_ *on the same taxon set*,

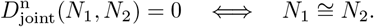

*Proof*. Suppose that 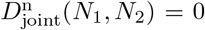, and let *M* be a minimizing matching. All terms in the objective are nonnegative, so every matched tripartition cost and the marked remainder cost are zero. Since matching a real event to a padding symbol has positive cost, *M* induces a bijection between *R*(*N*_1_) and *R*(*N*_2_).

For every *h ∈ R*(*N*_1_), the zero tripartition cost implies

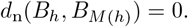

Binary level-1 networks are tree-child [8, Proposition 9], and this property is inherited by their descendant subnetworks. Thus, each *B*_*h*_ is tree-child. Since *d*_n_(*B*_*h*_, *B*_*M*(*h*)_) = 0, the two blocks have equal vertex-equivalence multiplicities and therefore the same taxon set. VED separates tree-child networks [3]; although the root *h* of *B*_*h*_ has outdegree one, their definition permits this and explicitly includes descendant subnetworks [3, Section 2.1]. Hence

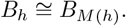

Likewise, the marked remainders are rooted trees whose leaf labels may repeat. VED separates such multi-labelled trees [3]. Their zero dissimilarity therefore gives

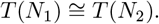

Lemma 6 now yields *N*_1_ *≅ N*_2_.

Conversely, an isomorphism *N*_1_ *≅ N*_2_ induces a reticulation matching under which corresponding tripartition blocks and marked remainders are isomorphic. Every term in the objective is therefore zero, so 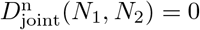. □

#### Theorem 3

(Identity of indiscernibles for the cluster instantiation). *Let N*_1_, *N*_2_ *be rooted binary level-*1 *networks on the same taxon set. Under the cluster instantiation*,

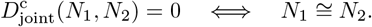

*Proof*. Suppose that 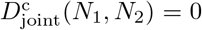. Let *M* be a minimizing matching. *M* induces a bijection between the reticulations. Corresponding hybrid blocks in reticulation tripartitions have equal cluster multisets, as do the marked remainders.

Let *ℋ*_*i*_ be the reticulations having no proper reticulation ancestor in *N*_*i*_. Equality of corresponding middle-block cluster multisets implies ℒ (*B*_*h*_) = ℒ (*B*_*M*(*h*)_) for every reticulation *h*. By Lemma 5, these sets distinguish reticulations within each network. Since the marked remainder cluster multisets contain the same marker labels, *M* (*ℋ*_1_) = *ℋ*_2_.

Recover the cluster multiset of *N*_*i*_ as follows. For each marker label, remove the two singleton cluster occurrences contributed by its marker leaves. In each remaining cluster, replace marker labels by the taxon sets they represent and take the union. Finally, add the cluster multisets of *B*_*h*_ for all *h ∈ ℋ*_*i*_. By Lemma 4, these blocks partition the removed vertices. This procedure recovers every original vertex’s cluster exactly once. Since each *B*_*h*_ contains all descendants of its vertices, their clusters within *B*_*h*_ are the same as in *N*_*i*_. Thus, this procedure recovers the full cluster multiset of *N*_*i*_.

The equal remainder and corresponding block cluster multisets yield equal cluster multisets for *N*_1_ and *N*_2_. Binary level-1 networks are determined by their cluster multisets [8, Corollary 44]; therefore, *N*_1_ *≅ N*_2_.

Conversely, an isomorphism *N*_1_ *≅ N*_2_ induces a reticulation matching under which corresponding tripartition blocks and marked remainders have equal cluster multisets. Hence 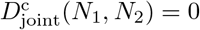. □

#### Corollary 1

(Metricity of the cluster and vertex-equivalence joint dissimilarities). *For each α ∈ {*c, n*}*, 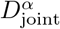 *is a metric on the class of rooted binary level-*1 *networks on a fixed taxon set*.

*Proof*. The joint dissimilarities are pseudometrics by Theorem 1. The preceding separation theorems establish identity of indiscernibles on rooted binary level-1 networks. Hence the cluster and vertex-equivalence instantiations are metrics on this class. □

#### Remark 1

(Failure of separation for the leaf-set instantiation). Let *T*_1_, *T*_2_ be non-isomorphic rooted binary trees on the same taxon set. Their reticulation dissimilarity is zero because neither tree has reticulations. Their remainders are the trees themselves and have identical leaf sets, so their remainder dissimilarity is also zero. Thus,

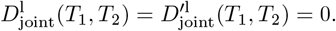

Neither the unnormalized nor the normalized leaf-set joint dissimilarity is therefore a metric, even on binary level-1 networks.

## 5 Empirical evaluation

We compare our joint dissimilarity with normalized hardwired-cluster distance (HWCD), normalized VED, and HAE; implementations and normalizations are specified in Appendix C. Our mathematical guarantees apply to unnormalized values, whereas experiments use normalized scores to compare trends on a common scale. We primarily show the vertex-equivalence instantiation; additional comparisons appear in the appendix. Reporting both 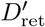 and *D*_tree_ allows both sources of disagreement to be assessed independently of their weights in the joint score.

### 5.4 Sensitivity to reticulation errors as the number of taxa grows

Standard normalized measures of overall network topology can assign small dissimilarities to networks and their displayed trees when the tree-like portions are extensive. Our dissimilarity explicitly penalizes missing reticulations by construction. This experiment illustrates how that penalty affects comparisons between networks and their displayed trees as the number of taxa increases.

For each taxon-set size *n ∈ {*50, 250, 500, 1000}, we simulated 100 networks, with 20 at each of five reticulation counts: 1,2,3,4,5 for 50 taxa; 1,3,5,7,10 for 250 taxa; 1,4,7,11,15 for 500 taxa; and 1,7, 13, 19, 25 for 1000 taxa. For each network, we sampled up to 32 distinct displayed trees. Because a displayed tree contains no reticulations, every reticulation in the network is unmatched, so the normalized reticulation term equals 1 by construction.

Figure 2 reports dissimilarities between the displayed trees and their corresponding networks under the different measures. For a single missing reticulation, the mean joint dissimilarity decreases modestly from 0.62 at 50 taxa to 0.51 at 1,000 taxa. The mean HWCD and normalized VED decrease from 0.09 and 0.11, respectively, at 50 taxa to 0.01 at 1,000 taxa. When comparing networks with five missing reticulations at 50 taxa against networks with 25 missing reticulations at 1,000 taxa, the mean HWCD and normalized VED decrease from 0.34 and 0.27 to 0.15 and 0.12, respectively. These results illustrate how the explicit penalty for missing reticulations maintains the joint dissimilarity’s sensitivity as network size grows, whereas the two global measures assign smaller values even when more reticulations are missing.

**Figure 2.**
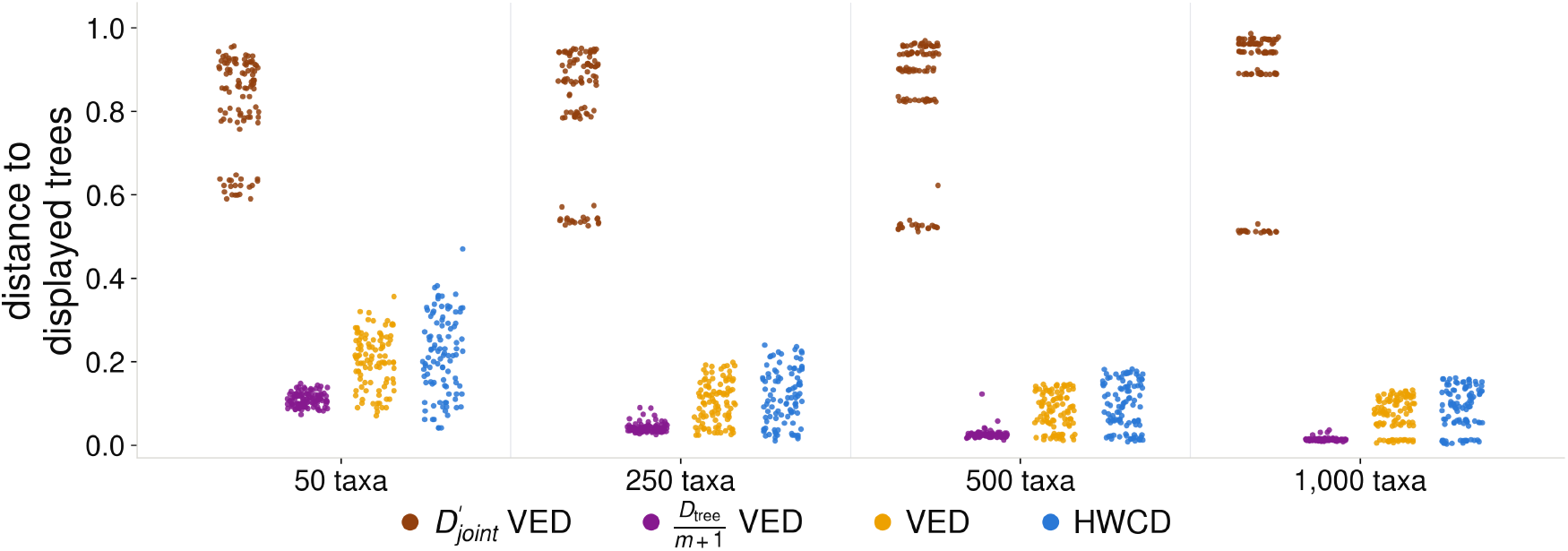
Measuring the dissimilarity between a network and its displayed tree with the proposed dissimilarity measure, VED, and HWCD. Each dot represents one network’s mean dissimilarity to its displayed trees. From left to right, each group shows the normalized joint dissimilarity (VED instantiation), its tree-remainder contribution *D*_tree_*/*(*m* + 1), normalized VED, and HWCD. Each taxon-set size has 100 reference networks, 20 at each reticulation count. HAE is omitted because it equals 0.5 under the empty-set convention for tree-only estimates.

When we consider only the tree-remainder contribution, *D*_tree_*/*(*m* + 1), displayed trees are close to the networks that display them, and this contribution becomes smaller as the number of taxa increases (Figure 2). Thus, even when a displayed tree has a large joint dissimilarity to its network because it lacks all of its reticulations, the tree remainder contributes little to that score.

We also examine the converse setting, in which the reference is a tree and the network contains an added reticulation (Appendix D.1). The mean tree-remainder contribution remained small, decreasing from 0.129 at 50 taxa to 0.013 at 1,000 taxa. Moving the spurious reticulation’s parent or hybrid attachment by one to three hops had little effect on the mean tree-remainder contribution.

### 5.2 Fixed reticulation error and perturbed backbones

We next examine how the dissimilarities respond to fixed reticulation errors as the tree-like portion grows and when backbone errors are introduced. We designed six small fixed networks with 12 to 25 taxa and constructed perturbations of these networks using wrong-hybrid or wrong-parent edits. A small network *N* is attached to a larger tree backbone *T* (Figure 3(a)). This construction allows us to vary small-network error and tree-part error separately.

**Figure 3.**
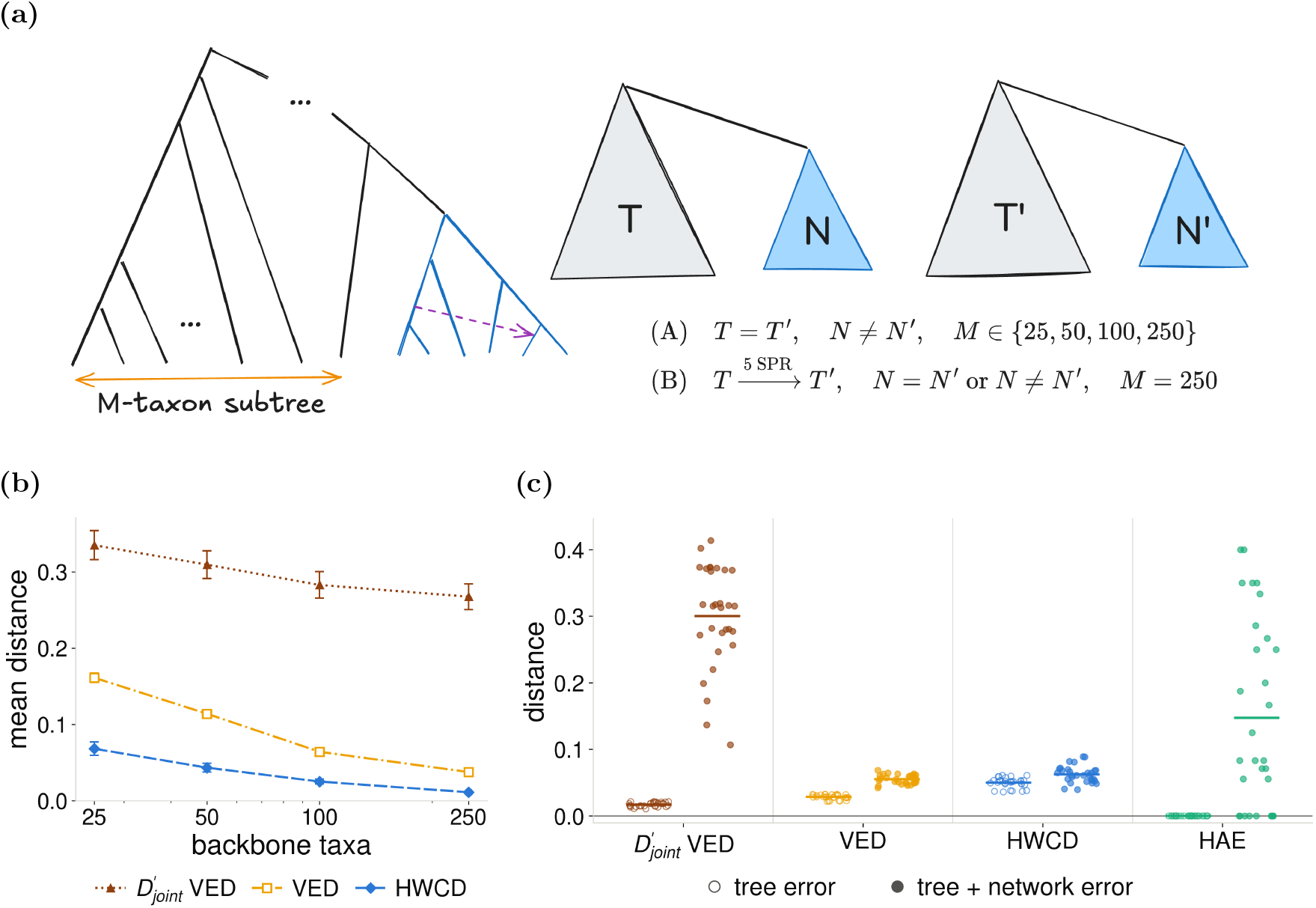
The change in dissimilarity when a small network attached to a bigger tree is perturbed, using the proposed dissimilarity measure, VED, HWCD, and HAE. **(a)** A small network (blue) is attached to a larger tree backbone. **(b)** Sensitivity to fixed small-network error as the correct backbone grows. Each small-network perturbation is compared with its reference within identical backbones of 25, 50, 100, or 250 taxa. Curves show the mean dissimilarity over the 30 comparisons at each backbone size (six small networks, five perturbations); error bars show one standard error of the mean. **(c)** Separating network error from tree-part error at 250 backbone taxa. One small network with two or three reticulations is embedded in a 250-taxon backbone tree. The backbone of each estimate is perturbed by five SPR moves. Hollow circle/left column: the small network is correct. Filled circle/right column: the small network is additionally perturbed by one random reticulation edit. Points show individual comparisons between the perturbed network and the truth, and horizontal bars show group means. Panel (c) shows our joint dissimilarity with the vertex-equivalence instantiation, the normalized VED, HWCD, and HAE.

We evaluated two settings: (i) we perturbed the small network while keeping the tree backbone correct, increasing the number of backbone taxa from 25 to 50, 100, and 250; (ii) we fixed the backbone at 250 taxa and applied five SPR moves, then compared estimates containing either the unchanged or perturbed small network on that same perturbed backbone. In the first setting, the reticulation contribution is fixed by construction. We examine how this contribution affects the joint score as the correct tree-like portion grows, relative to the other measures. The second setting examines how the measures respond to additional reticulation error when both estimates share the same perturbed backbone (Figure 3(c)).

In the first setting, all three measures decrease as the number of backbone taxa increases, but by different amounts (Figure 3(b)). The normalized VED decreases from 0.161 to 0.038, a reduction by a factor of approximately 4.2, while HWCD decreases by a factor of 6.2 to 0.011. Nevertheless, the joint dissimilarity decreases only from 0.34 to 0.27. Its two components behave differently: the reticulation contribution to the normalized joint score remains constant at 0.25, while the tree-remainder contribution decreases from to 0.02. These results illustrate how the explicit reticulation contribution maintains sensitivity to the fixed error as the correct tree-part grows. In the second setting, estimates containing the perturbed small network receive higher joint dissimilarity scores than those containing the unchanged small network on the same perturbed backbone. The score difference is larger than under the normalized VED or HWCD in these comparisons. HAE detects some reticulation perturbations that alter hybrid ancestry but remains insensitive to wrong-parent edits. Additional examples of structurally distinct networks with zero HAE are presented in the appendix (Figure 10).

We emphasize that the constant reticulation contribution in this backbone experiment, rather than the absolute joint value in Figure 2, is the clearer evidence of scale-robust sensitivity. In the displayed-tree comparison, every reticulation is unmatched by construction, so the normalized reticulation term equals one and, for a network with *m* reticulations, the normalized joint dissimilarity is bounded from below by *m/*(*m*+1), independently of the number of taxa. What Figure 2 helps establish, therefore, is the comparisons with several reticulations and the contrast with the global measures, which tend toward zero over the same range, rather than the magnitude of the joint value itself.

### 5.3 Distinguishing errors in tree-of-blobs resolution

A tree of blobs (TOB) summarizes the relationships among reticulated regions by contracting each blob to a single vertex. Resolving a blob requires specifying its internal network structure. Figure 4(a) shows a resolved blob with descendant groups *g*_0_ through *g*_5_ and illustrates four possible errors in its resolution: (i) the hybrid group is wrongly swapped with a parental group; (ii) the hybrid group is wrong while both parental groups are correct; (iii) one parental group is wrong while the hybrid is correct; (iv) two internal groups are wrongly swapped while both the hybrid and the parents are correct.

**Figure 4.**
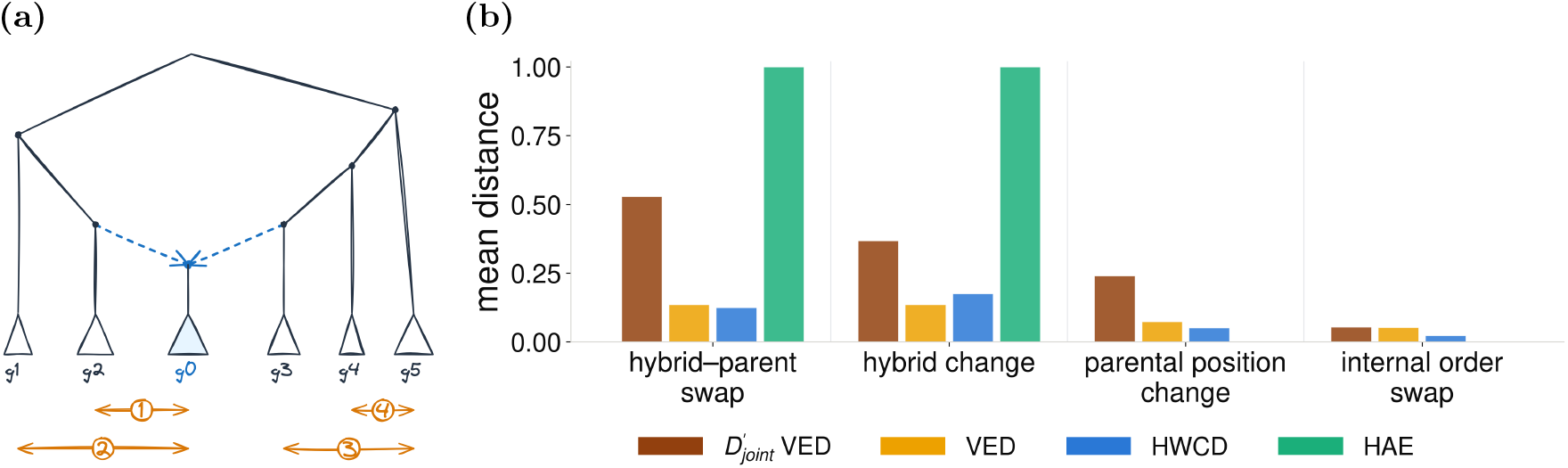
Joint dissimilarity, VED, HWCD, and HAE between the networks under four types of error in tree-of-blobs resolution. **(a)** Triangles represent taxon groups; *g*_0_ is the hybrid and *g*_2_, *g*_3_ are parental groups. Orange arrows 1–4 indicate errors that preserve the tree of blobs. **(b)** Dissimilarities for the four error categories, ordered from left to right as in (a). Bars are means over 20 comparisons per error category.

We simulated ten binary level-1 networks with 44 taxa, each containing one reticulation in a single blob with ten attached groups. These groups collectively contain 41 taxa, with the remaining three taxa forming an outside group. For each reference, we generated two distinct resolutions per error category, yielding 80 comparisons. Every perturbed network preserves the reference tree of blobs, so the comparisons isolate differences in how its single blob is resolved.

The dissimilarities between the perturbed networks and the ground-truth networks are shown in Figure 4(b). The joint dissimilarity separates all four categories, whereas the normalized VED conflates hybrid-parent swaps with hybrid changes, HWCD ranks hybrid changes above hybrid-parent swaps, the reverse of our ordering, and HAE is sensitive only to changes in hybrid identity. The relative ordering of the four categories under our measure follows from how the tripartition construction weights hybrid identity, parental placement, and internal structure, and is not claimed to be a ground-truth ranking of error severity. The point of this experiment is that these error types are separable under a single measure, not that one is objectively worse than another.

### 5.4 Calibration study

We assessed responses to cumulative errors and to the displacement of individual reticulations. We measure displacement in hops: hop distance is the minimum number of transitions between branches sharing a node, from the original attachment branch to the target branch, which is subdivided for reattachment.

#### Number of changed reticulations

We generated 40 reference networks, each with five reticulations: 20 with 50 taxa and 20 with 100 taxa. For each network, we constructed a sequence of perturbed networks, adding one wrong-parent or wrong-hybrid edit at each step. Each new network retained the preceding edits, with the new attachment selected from branches one to three hops away. A step was accepted only when exactly *j* reticulations differed from the reference. We retained only networks reaching *j* = 5: 33 networks for wrong-parent edits and all 40 for wrong-hybrid edits. Thus, every point within each panel averages over the same reference networks.

Figure 5 shows approximately linear increases in dissimilarity with the number of changed reticulations. The joint dissimilarity reaches 0.31 for five wrong-parent edits (Figure 5(a)) and 0.64 for five wrong-hybrid edits (Figure 5(b)). The normalized VED and HWCD also increase, reaching 0.16 and 0.13, respectively, for wrong-parent edits and 0.28 and 0.32 for wrong-hybrid edits. HAE remains zero under wrong-parent edits but increases to 0.55 under wrong-hybrid edits. The joint dissimilarity therefore responds to increasing reticulation error in both families, with wrong-hybrid edits producing approximately twice the mean dissimilarity of wrong-parent edits.

**Figure 5.**
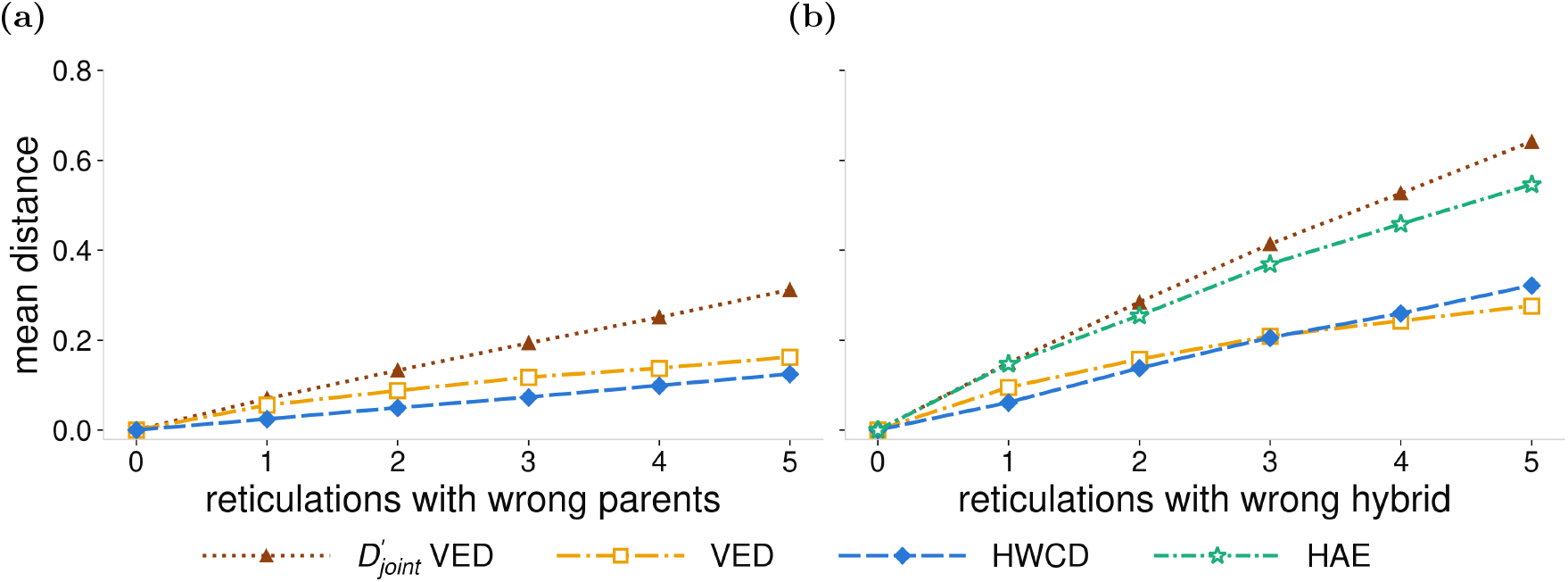
Change in the joint dissimilarity, VED, HWCD, and HAE as the number of incorrect reticulations increases under (a) wrong-parent edits and (b) wrong-hybrid edits. Each reference has five reticulations and 50 or 100 taxa. Every point averages over the same 33 wrong-parent or 40 wrong-hybrid networks; new attachments are one to three hops away. HAE is identically zero for wrong-parent edits and is omitted in (a).

#### Magnitude of an individual error

To distinguish the number of errors from their magnitude, we generated 30 reference networks with 100 taxa and three reticulations. We moved one hybrid or one parental attachment to a target branch exactly one, three, or six hops away; for wrong-parent edits, the same incoming reticulation edge was selected at each distance.

The joint dissimilarity is the only measure whose median increases at every displacement level under both edit families (Figure 6(a), (b)): from 0.20 to 0.22 to 0.23 for wrong-hybrid edits, and from 0.090 to 0.102 to 0.107 for wrong-parent edits. Normalized VED has identical medians at one and three hops, followed by a small increase at six hops: 0.071, 0.071, and 0.078 for wrong-hybrid edits, and 0.039, 0.039, and 0.044 for wrong-parent edits. HWCD has a constant median of 0.067 under wrong-hybrid edits but increases from 0.010 to 0.058 under wrong-parent edits. HAE increases from 0.13 to 0.43 under wrong-hybrid edits and remains zero under wrong-parent edits.

**Figure 6.**
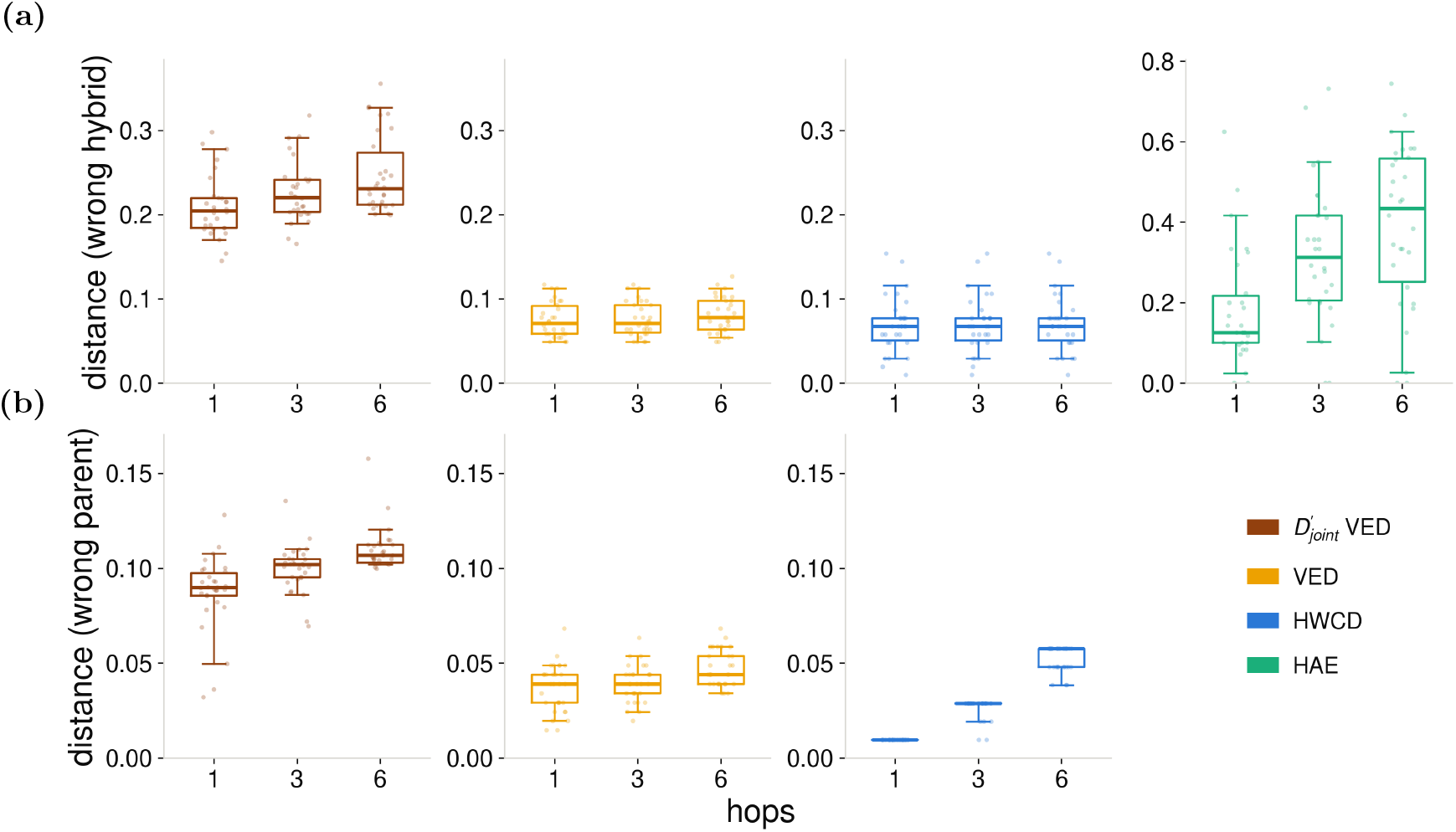
Changes in dissimilarity as a single reticulation has increasing magnitude of displacement. On networks with 100 taxa and three reticulations, **(a)** wrong-hybrid edits and **(b)** wrong-parent edits are applied to one reticulation. Columns show the joint dissimilarity, VED, HWCD, and HAE from left to right. HAE is zero for wrong-parent edits and is omitted from the bottom row. Boxes show interquartile ranges, lines medians, whiskers the 5th–95th percentiles, and dots individual networks. The first three panels in each row share a scale; HAE has its own.

We performed a separate backbone calibration which keeps reticulations correct while increasing tree-part error. The reticulation term remains zero, whereas the mean remainder term increases as the RF distance in the tree part increases (Appendix D.3). Together, these controlled settings illustrate complementary responses to reticulation and backbone disagreement.

### 5.5 Assessment of inferred networks under different dissimilarity measures

We evaluated ASTRAL [26], CAMUS [25], NetCS [5], InPhyNet [12], DIMPLE [7], and BROOQS [2] on 30 simulated networks with 50 taxa under low incomplete lineage sorting (ILS): ten each of level 1, level 2, and level 3 or higher. These networks came from the DIMPLE simulations [7] and were generated under a birth–death–hybridization model using SiPhyNetwork [10].

Figure 7 shows the resulting dissimilarities, with ASTRAL serving as a tree-only control. All measures assign the lowest mean dissimilarity to DIMPLE. Under the joint dissimilarity, its mean is 0.33, compared with 0.50–0.58 for the other network inference methods and 0.78 for ASTRAL.

**Figure 7.**
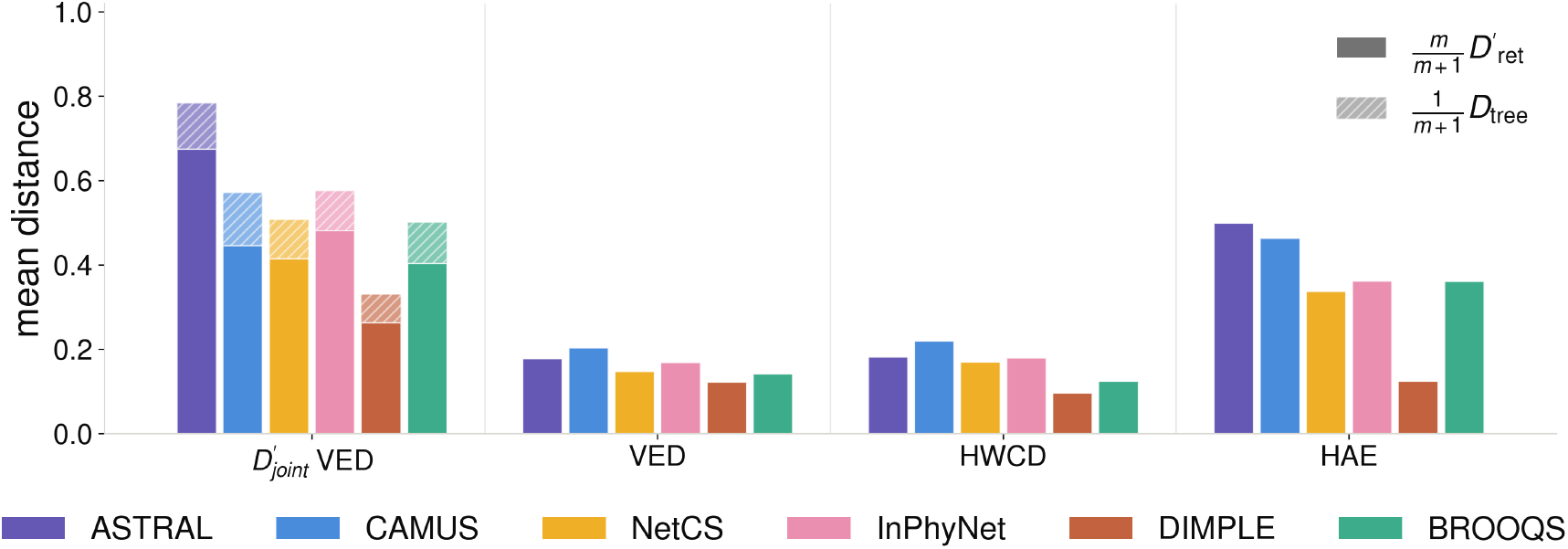
Reporting dissimilarities between the true network and networks inferred by different methods using the joint dissimilarity, VED, HWCD, and HAE. Bars show mean normalized dissimilarities on 30 networks. For the joint score, opaque and hatched segments show the weighted reticulation and remainder contributions, respectively.

The measures differ in their assessment of the tree-only control. The normalized VED and HWCD both assign ASTRAL a mean score of 0.18—lower than CAMUS’s (0.20 and 0.22) and comparable to InPhyNet’s (0.17 and 0.18)—even though ASTRAL’s estimates lack all reticulations in the reference networks. This is consistent with the limited sensitivity to missing reticulations observed in Figure 2.

In contrast, the joint dissimilarity assigns ASTRAL the largest mean score. Examining the two terms separately clarifies this ranking. ASTRAL’s normalized reticulation term is saturated at 1 because its tree estimates contain no reticulations. Its mean tree-remainder term is 0.37, lower than CAMUS’s 0.45. CAMUS therefore has less reticulation disagreement but greater tree-remainder disagreement than ASTRAL. The decomposition reveals these contrasting sources of error that are obscured by a single overall score.

### 5.6 Runtime

Our joint dissimilarity is fast to compute: the vertex-equivalence instantiation in PhyNetPy has a median CPU time of 54.4 ms per network pair at 1,000 taxa, across networks with up to 25 reticulations. Detailed runtime comparisons are provided in Appendix D.5 (Figure 13).

## 6 Discussion

In this work, we introduce a family of network dissimilarities that explicitly compare reticulation structure through optimal matching of reticulation tripartitions. Each tripartition captures the hybrid and parental subnetworks of a reticulation. Combining this comparison with a tree-remainder term, computed after removing the hybrid components, yields a pseudometric on rooted binary phylogenetic networks on the same taxon set. We develop three instantiations, of which the cluster and the vertex-equivalence instantiations are metrics on binary level-1 phylogenetic networks. Computationally, our construction permits polynomial-time computation using a single assignment solve and one remainder comparison.

Experiments show that our measures clearly separate trees and networks and retain sensitivity to missing and misplaced reticulations even when the tree-like portion of the network is large. While HWCD and the normalized VED can approach zero for a fixed reticulation error as the number of taxa in the tree-part increases, our joint dissimilarity remains sensitive to that error and maintains variation in scores across networks with different numbers of missing reticulations, preserving their separation within each dataset size. These results show that a small overall dissimilarity can conceal substantial reticulation error when most of the tree-like portion of the network is correct. Explicitly retaining the reticulation contribution helps prevent agreement in the tree backbone from dominating the assessment of network reconstruction.

In our experiments, our dissimilarities assign larger penalties to errors in hybrid identity than to errors in parental placement, as observed in Figures 5, 6, and 4(b). Distinguishing hybrid identity, parental placement, and internal ordering makes the measures useful for evaluating methods that resolve trees of blobs into level-1 networks. Beyond distinguishing error types, the measures show graded responses to both the number of incorrect reticulations and the magnitude of their displacement.

HWCD is less responsive to some changes, as it counts unmatched clusters rather than the magnitude of disagreement within those clusters. Different displacements can therefore receive identical scores when they produce the same number of unmatched clusters. Similarly, in our local NNI experiment (Appendix, Figure 12), HWCD remains unchanged with increasing edit depth because each rearrangement replaces one cluster with another, regardless of its depth. The vertex-equivalence instantiation responds to depth because the change affects the structural representations of the affected vertex and its ancestors. These results highlight differences in what the measures quantify beyond metricity alone.

HAE provides useful complementary information about hybrid ancestry, separating some perturbed networks and identifying the limitations of tree-only estimates. However, it misses important structural changes that preserve hybrid ancestry. In particular, it does not detect any errors in parental placement that leave hybrid ancestry unchanged. In our tree-of-blobs experiment (Section 5.3), HAE is zero for both parental-position changes and internal-order swaps, despite the resulting networks being incorrect. Comparisons based on HAE alone can therefore fail to penalize networks with incorrect parental placement. This motivates evaluating hybrid ancestry alongside structural agreement.

The tree-remainder term helps distinguish disagreement in the tree-like portion of the network from disagreement about its reticulations. In Figure 2, each displayed tree lacks all of its network’s reticulations, yet the tree-remainder contribution to the joint score is small and decreases on average as the number of taxa grows. With one spurious reticulation added to each ground truth tree (Appendix D.1), the mean tree-remainder contribution was small and changed little when we moved the reticulation, reflecting agreement with the true tree despite the incorrect reticulation. This distinction also helps interpret inferred networks. On average, ASTRAL has a larger reticulation term than CAMUS but a smaller tree-remainder term, showing that their overall scores reflect different combinations of error. We therefore recommend reporting the reticulation and tree-remainder terms alongside the joint score.

Several limitations remain and motivate further work. We prove that the unnormalized cluster and vertex-equivalence instantiations are metrics on binary level-1 networks. None of the three joint dissimilarities is a metric on all rooted binary networks, as demonstrated by the level-6 counterexample in Appendix A (Figure 8). Determining which broader network classes are distinguished by these representations, including tree-child networks, and modifying the representations where necessary to separate networks in more general classes remain important directions. Second, missing and incorrect reticulations can receive equal penalties under the reticulation term, although they represent different failures of reconstruction. Separately reporting unmatched reticulations and disagreement between matched reticulations could make this distinction clearer. Examining these contributions alongside the tree-remainder term, and investigating alternative relative weights, could further improve the interpretation of inferred-network comparisons. Finally, the reticulation-matching component is analogous to pairwise sequence alignment: matched reticulations incur a cost for structural disagreement, while unmatched reticulations incur a penalty similar to a gap cost. This analogy suggests a natural extension to multiple-network comparison, which could help identify corresponding reticulations across several inferred networks and summarize agreement and uncertainty in their reticulate histories.

**Figure 8.**
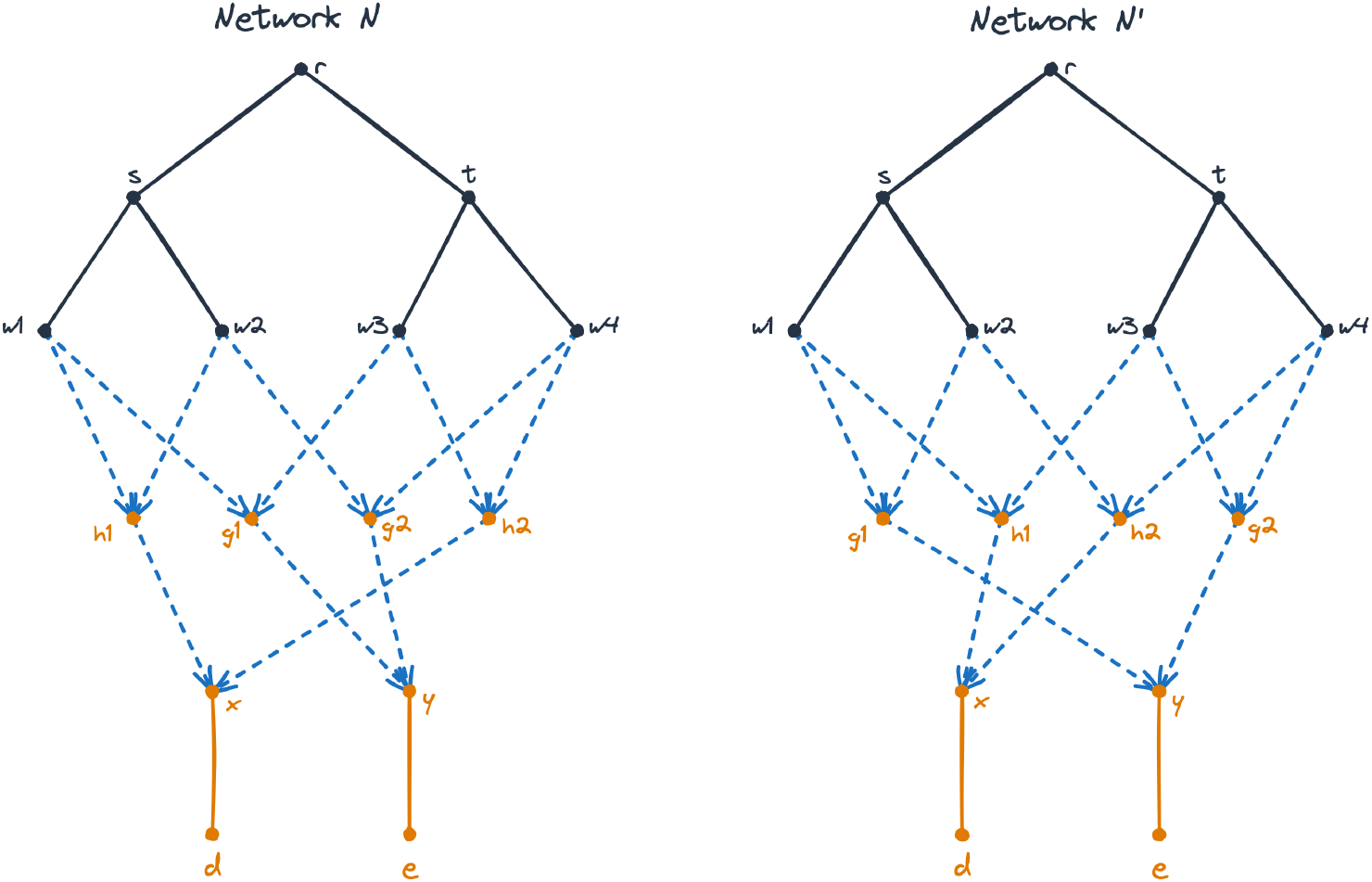
Non-isomorphic binary level-6 networks with zero joint dissimilarity under all three instantiations. Only *d* and *e* are taxon labels; internal vertex names are included for reference. Dashed blue arrows indicate reticulation edges.

## Data Availability

The dissimilarities are implemented in the PhyNetPy library, available at https://github.com/NakhlehLab/PhyNetPy. All the data simulated for the study is available at https://doi.org/10.5281/zenodo.23933421.

## Declaration of AI-assisted technologies

During the preparation of this work, the authors used OpenAI’s Codex (GPT-6 Astra) and Anthropic’s Claude (Fable 5) to assist with editing manuscript text, developing proofs and analysis code, and preparing figures. The authors reviewed and edited the resulting material, verified the analyses, and take full responsibility for the content of this article.

## APPENDIX

### A Failure of separation beyond level-1 networks

Figure 8 shows two non-isomorphic binary level-6 networks on {*d, e}* with zero joint dissimilarity under all three instantiations. To see that they are not isomorphic: in *N* , the two parents of each parent of *x* share a parent: *w*_1_, *w*_2_ share *s*, and *w*_3_, *w*_4_ share *t*. In *N*^*′*^, the corresponding pairs *w*_1_, *w*_3_ and *w*_2_, *w*_4_ do not share a parent.

Matching equally named reticulations gives isomorphic middle blocks and isomorphic parental blocks. Moreover, each *w*_*i*_ has two marker leaves in the remainder, labelled by *{d}* and *{e}*, so the marked remainders are isomorphic. Therefore, for every *α ∈ {*l, c, n*}*,

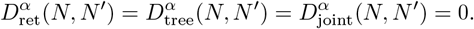

Thus, none of the three joint dissimilarities is a metric on all rooted binary phylogenetic networks.

### B Algorithm for computing the joint dissimilarity

#### Algorithm 1 Computation of the joint dissimilarity

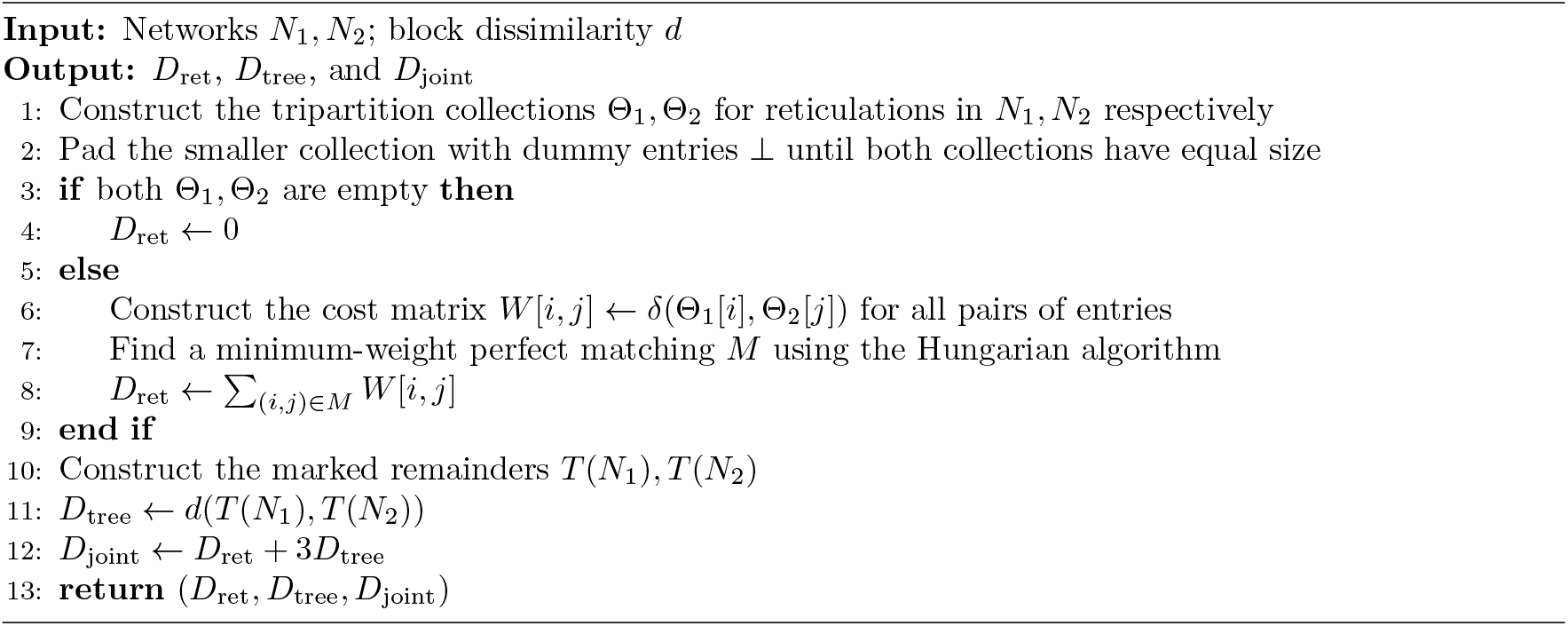

### C Comparison measures

HWCD was computed following PhyloNetworks.jl’s hardwiredclusterdistance function [24]. We divide the total number of unmatched occurrences across both multisets of clusters by their combined size. We computed VED [21] with the `Nakhleh_metric` function in PhyNetPy 0.7.0 [11] and normalized it as 2*D/*(|*V*_1_| + |*V*_2_|), where *D* is the raw distance and *V*_1_, *V*_2_ are the vertex sets.

HAE [5] was computed using the `hybrid_ancestry_error` function in PhyNetPy. It compares the sets of taxa with hybrid ancestry in the true and estimated networks. It averages the proportion of the estimate’s hybrid-ancestry taxa absent from the truth’s set and the proportion of the truth’s hybrid-ancestry taxa absent from the estimate’s set. Any error proportion with a zero denominator is defined to be zero. All three measures take values in [0, 1].

### D Additional experiments

#### D.1 Tree-remainder contribution

We used 80 displayed trees from Section 5.1 as ground truth trees: 20 each at 50, 250, 500, and 1,000 taxa. We added one reticulation to each tree and then constructed two variants, moving either the parent attachment or the hybrid attachment by one to three hops. All three networks displayed the original tree.

The mean tree-remainder contribution decreased from 0.129 at 50 taxa to 0.013 at 1,000 taxa for the added reticulations (Figure 9). Moving the attachment had little effect on this mean: the mean differed from that for the initial placement by less than 0.004 at every dataset size.

**Figure 9.**
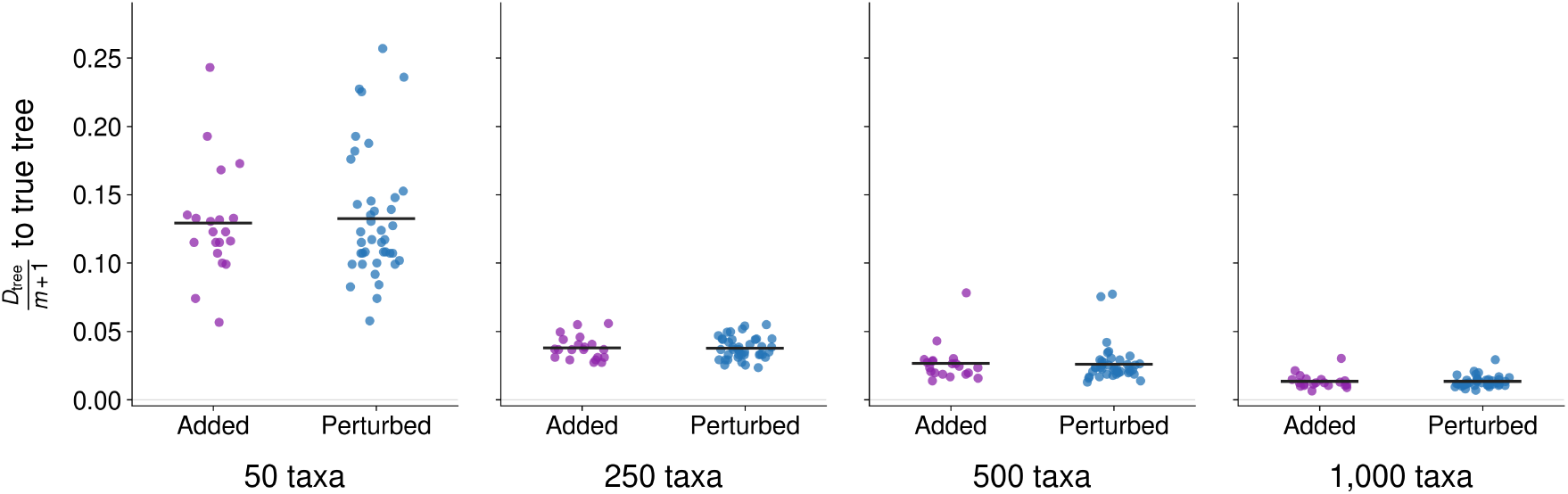
Measuring tree-remainder contribution to the joint dissimilarity (VED instantiation) between a reference tree and a network that displays it. *Added* : one reticulation is added to the tree (20 comparisons per dataset size). *Perturbed* : the same reticulation is then moved (one parent move and one hybrid move per tree; 40 comparisons). Each network has one reticulation. Dots show individual comparisons; horizontal bars show means.

#### D.2 Structural differences undetected by HAE

We highlighted examples of structural differences that cannot be detected by HAE in Figure 10. These networks have the same set of taxa with hybrid ancestry but differ in parental placement, reticulation structure, or the topology below a reticulation.

**Figure 10.**
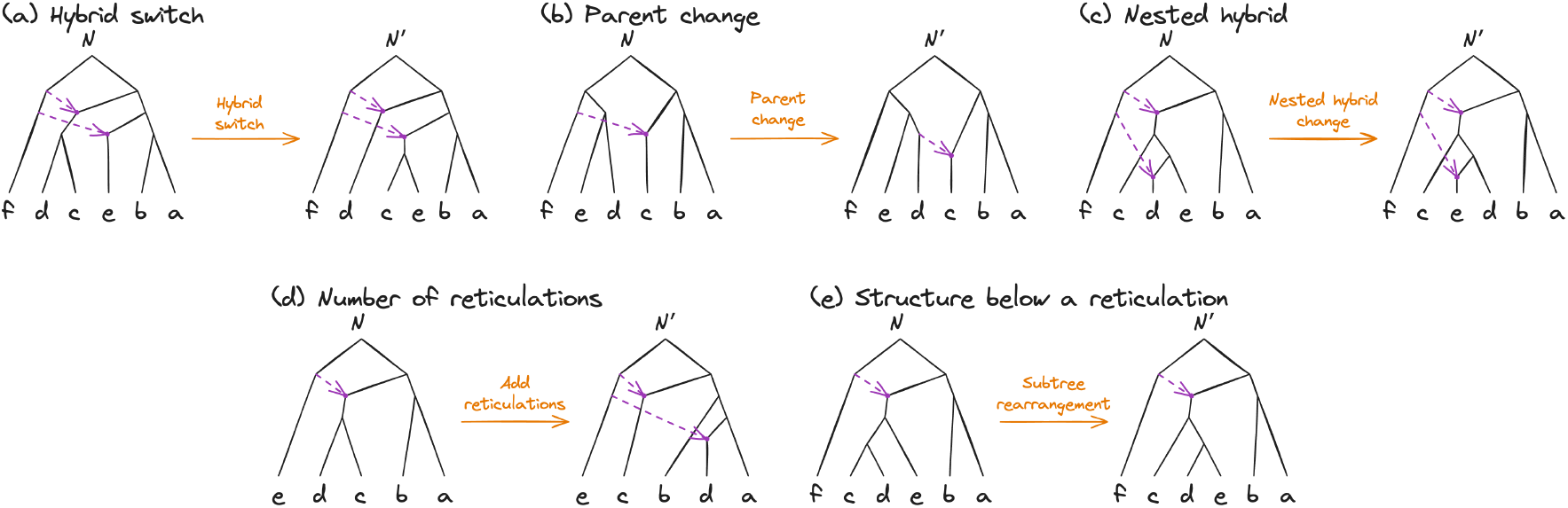
Examples of structural differences invisible to HAE. Each pair shows networks *N* and *N* ^*′*^ with different topologies but the same set of taxa with hybrid ancestry, giving HAE equal to zero. Purple dashed arrows mark reticulation edges, and orange arrows describe the changes: (a) switching hybrid placement; (b) changing a parent; (c) changing a nested reticulation; (d) adding a reticulation without changing hybrid ancestry for additional taxa; and (e) rearranging the subtree below a reticulation. These examples illustrate that HAE does not distinguish reticulations that preserve the set of taxa with hybrid ancestry.

#### D.3 Calibration against tree-part error

Figure 11 examines how dissimilarity responds to increasing tree-part error when the reticulations remain correct. Each of six small networks with two to four reticulations is grafted into five backbone trees with 100 taxa. We perturb only the backbone using random SPR moves, accepting rearrangements whose normalized rooted Robinson–Foulds (RF) distance from the original backbone falls in one of three ranges: [0.05, 0.15], [0.25, 0.35], or [0.55, 0.65]. For each small-network–backbone combination, we retain five samples per range.

**Figure 11.**
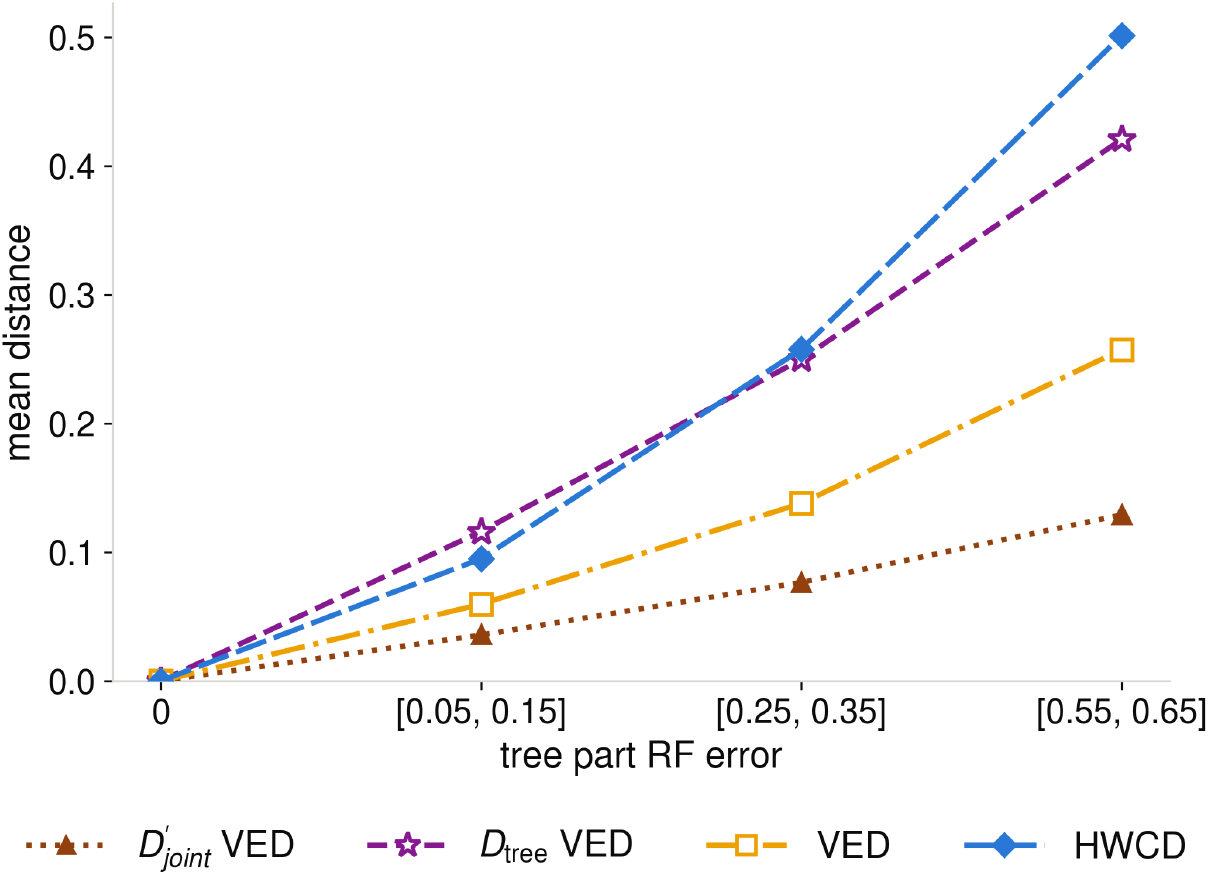
Calibration of joint dissimilarity, *D*_tree_, VED, and HWCD against increasing tree-part error when reticulations stay correct. Six small networks are each embedded in five trees with 100 taxa. SPR perturbations are applied to the tree part and summarized by normalized rooted RF ranges. Each point at a nonzero RF range shows the mean over 150 comparisons. The reticulation dissimilarity and HAE are both zero.

The reticulation term is zero in every comparison. The mean remainder term *D*_tree_ increases from 0.116 to 0.250 to 0.421 as the RF error range increases. HWCD and the normalized VED also increase across the RF ranges, from 0.095 to 0.258 to 0.501 and from 0.060 to 0.138 to 0.257, respectively. Together with Figure 3, this experiment illustrates how the two terms separate reticulation error from tree-part error under these controlled perturbations.

#### D.2 Differences between cluster and vertex-equivalence instantiations

For this experiment, we illustrate one difference between cluster and vertex-equivalence instantiations of our joint distances. Figure 12(a), right, illustrates the rationale in a tree: an NNI changes only the cluster of *x*, regardless of depth, but changes the vertex-equivalence classes of *x* and its ancestors — *d* + 2 vertices when *v* has depth *d*. The experiment thus tests whether dissimilarity is independent of the position of a change or increases with its depth.

We generated 20 random rooted binary trees at each of 50, 100 and 250 taxa, then added three reticulations to each tree. We define the depth of a vertex as the number of edges on the shortest directed path from the root to it. At every depth *d* = 0, … , 12, we selected one vertex uniformly at random and applied a single nearest-neighbour interchange (NNI). As shown in Figure 12(a), left, the NNI exchanges subtrees *y* and *q*, where *v* has children *x* and *y*, and *x* has children *p* and *q*. Both *v* and *x* must be tree vertices with two children; the exchanged subtrees may contain reticulations. We scored each comparison between an altered network and its original network using the joint distance under the cluster and vertex-equivalence instantiations, HWCD, and the VED.

**Figure 12.**
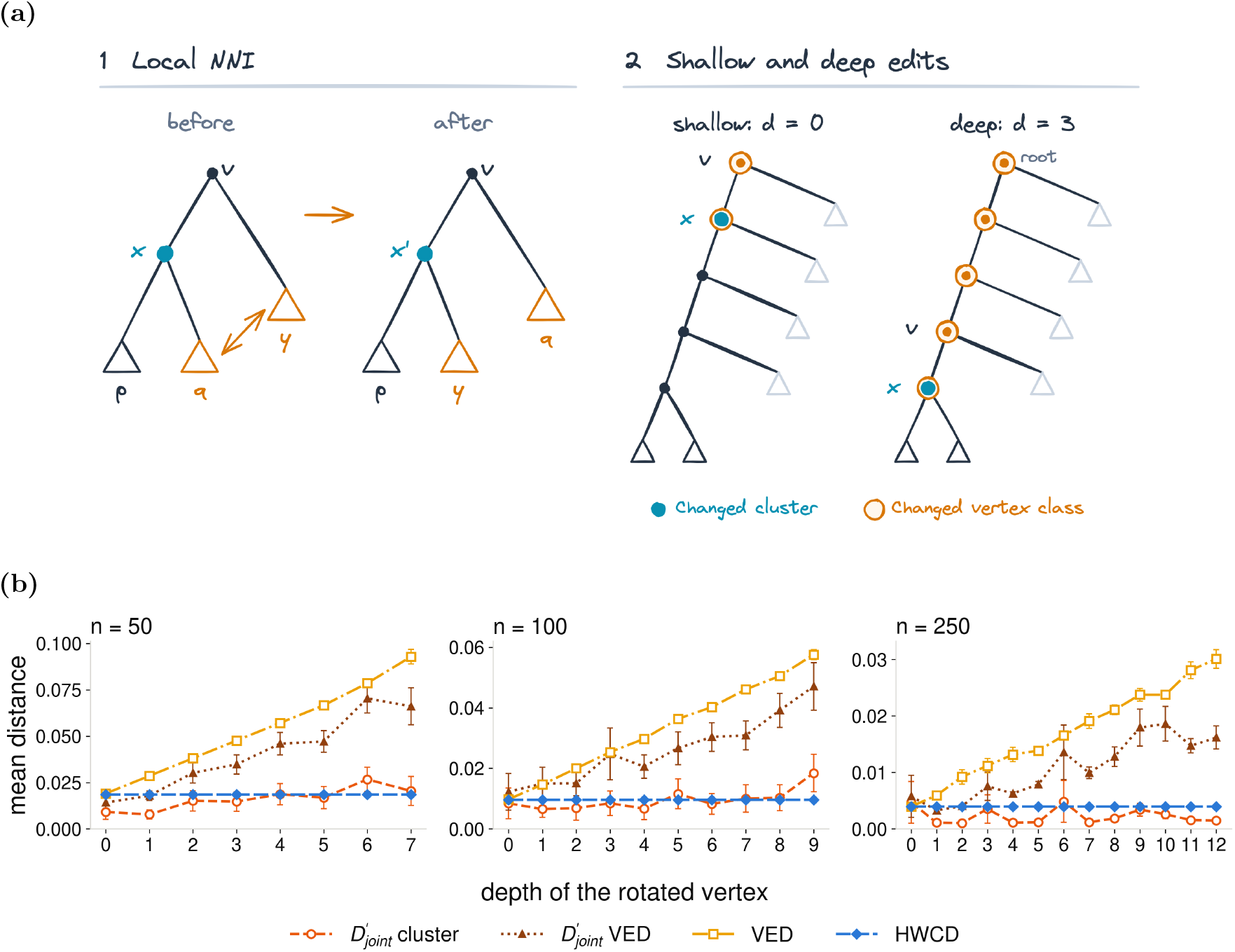
Measuring dissimilarities when the depth of error in the network increases. **(a)** Left: the local NNI shows the exchange of subtrees *y* and *q*. Right: shallow and deep edits in a tree are compared: blue marks the changed cluster at *x*, and orange marks the vertices whose vertex-equivalence classes change. **(b)** Dissimilarity versus NNI depth for networks with 50, 100 and 250 taxa and three reticulations. Depth is measured from the root to *v*. Points are means over the same 20 networks with standard-error bars. Curves show the joint distance under the cluster and vertex-equivalence instantiations, the VED, and HWCD.

The results in Figure 12(b) include only depths at which all 20 ground truth networks of a given taxon size had an eligible vertex: 0–7 for 50 taxa, 0–9 for 100 taxa, and 0–12 for 250 taxa. Each plotted point averages over the same 20 truths.

HWCD remains constant across depths, while the cluster-based joint dissimilarity fluctuates without a consistent increasing trend. This is because an NNI can change only the hardwired cluster of *x*; the vertex-equivalence instantiation and the VED grow with depth as the rotation re-labels the rotated vertex and every one of its ancestors. The oscillations in the curves mark depths at which more rotations fell inside a reticulation’s block and so increased the reticulation term of our joint distance.

#### D.5 Runtime comparison

We measured CPU time on 95 pairs of reference networks from the displayed-tree experiment at each of 50, 250, 500, and 1,000 taxa, with all measurements performed in a single job on the same compute node. Networks within each pair had the same number of reticulations, with up to 5, 10, 15, and 25 reticulations at the four dataset sizes, respectively. The joint dissimilarity, VED, and HAE were timed through PhyNetPy 0.7.0 and HWCD through PhyloNetworks.jl; parsing time is excluded.

The median CPU time for our joint dissimilarity increased from 2.1 ms at 50 taxa to 54.4 ms at 1,000 taxa (Figure 13). At 1,000 taxa, median times were 10.5 ms for HAE, 9.1 ms for VED, and 4.92 s for HWCD. Our joint dissimilarity remained fast across the tested dataset sizes and numbers of reticulations.

**Figure 13.**
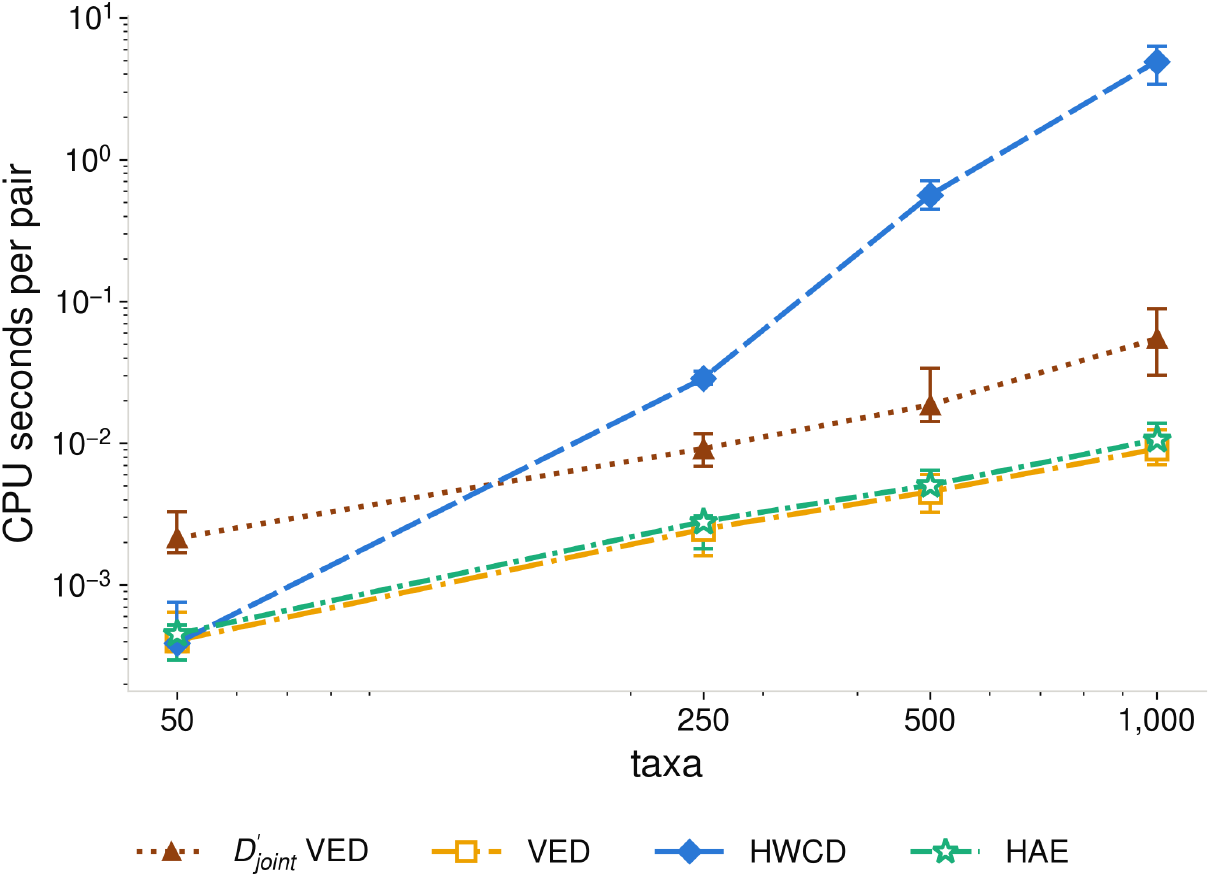
CPU time per network pair for the vertex-equivalence instantiation of the joint dissimilarity, normalized VED, HWCD, and HAE. Points show medians over 95 pairs at each dataset size; error bars span the 25th–75th percentiles. Both axes use logarithmic scales.

## Footnotes

1 The measure introduced here is completely different from those in [22, 18], despite the shared use of the term “tripartition.”

